# Integration of spatial and single nucleus transcriptomics to map gene expression in the developing mouse kidney

**DOI:** 10.1101/2024.12.18.629207

**Authors:** Christopher P. Chaney, Alexandria N. Fusco, Elyse D. Grilli, Jane N. Warshaw, Peter M. Luo, Ondine Cleaver, Denise K. Marciano, Thomas J. Carroll

## Abstract

The kidney is a complex organ requiring tightly coordinated interactions between epithelial, endothelial, and mesenchymal cells during development. Congenital kidney defects can result in kidney disease and renal failure, highlighting the importance of understanding kidney formation mechanisms. Advances in RNA sequencing have revealed remarkable cellular heterogeneity, especially in the kidney stroma, though relationships between stromal, epithelial, and endothelial cells remain unclear.

This study presents a comprehensive gene expression atlas of embryonic and postnatal kidneys, integrating single-nucleus and in situ RNA sequencing data. We developed the Kidney Spatial Transcriptome Analysis Tool (KSTAT), enabling researchers to identify cell locations, predict cell-cell communication, and map gene pathway activity. KSTAT revealed significant heterogeneity among embryonic kidney pericytes, providing a critical resource for hypothesis generation and advancing knowledge of kidney development and disease.

## Introduction

Recent advances in single-cell and single-nucleus RNA sequencing have revolutionized our understanding of cellular composition and signaling. These technologies have uncovered novel cell types, unexpected transcriptional states, and key insights into cellular differentiation. By simultaneously profiling all cells within a tissue, these approaches provide an unbiased view of cellular composition and behavior. This has led to a deeper understanding of relationships among cells, including transitions between cellular states during development, regeneration, and injury response. Rapid technological advancements now enable the analysis of tens of thousands of transcriptomes across various species, organs, and experimental conditions.

However, a critical aspect of cellular heterogeneity remains poorly understood: the spatial location of transcriptionally diverse cell states within their native tissue environment. Single-cell RNA sequencing relies on tissue dissociation for microfluidic capture, which results in the loss of spatial information. This limitation hinders our ability to reconstruct the complex cellular landscape and its dynamics accurately.

The spatial location of a cell determines how it interacts with neighboring cells, responds to biochemical cues, and accesses essential nutrients and oxygen from the bloodstream. Integrating spatial information with transcriptome data can yield novel insights into cellular behavior, interactions, and tissue organization. Spatial transcriptomics offers the potential to provide high-resolution information on tissue composition and function.

The kidney is a highly complex organ that plays a critical role in maintaining blood chemistry. It achieves this through functional units known as nephrons, epithelial tubules patterned along a proximal-distal axis into distinct functional domains. An adult human kidney contains, on average, one million nephrons per kidney. Defects in the patterning and differentiation of the kidney and ureter can impair renal function, potentially necessitating dialysis or transplantation to sustain life. Congenital anomalies of the kidney and urinary tract (CAKUT) account for approximately 10% of all identified birth defects.^1^ Studies across populations estimate the prevalence of CAKUT ranges from 1.6 to 20 cases per 1,000 births.^2,3^ Additionally, deficits in nephron number have been implicated in systemic hypertension^4^ and chronic kidney disease.^5^ A deeper understanding of kidney development is therefore essential for improving the diagnosis and treatment of birth defects and for addressing subtler contributions to systemic disease arising from aberrant kidney development.

The developing kidney is a complex organ composed of multiple cell types derived from at least four distinct developmental lineages, with intricate interactions between epithelial, endothelial, and mesenchymal cells.^6–8^ Recent single-cell RNA sequencing studies have highlighted molecular heterogeneity among embryonic renal interstitial cells,^9^ suggesting that these diverse populations create unique microenvironments that influence the differentiation of adjacent epithelial and endothelial cells.

The cellular complexity of the kidney poses significant challenges for researchers investigating its form and function. Analyses that lack spatial context risk misinterpretation of data, particularly in studies of cell-cell signaling. Recent efforts have aimed to create spatial atlases of gene expression in the embryonic kidney.^10–14^ While valuable, these resources are often constrained by the resolution of molecular localization technologies and the breadth or depth of gene coverage, meaning the number of genes detected or the number of transcripts detected per gene.

In this study, we address these limitations by combining single-cell transcriptomics with high-resolution spatial transcriptomics to generate a virtual atlas of embryonic and early postnatal mouse kidney tissue. Our analysis emphasizes the recently characterized heterogeneous renal stroma stroma,^9,15^ accurately predicting the expression and activity of genes and pathways beyond the initial set of landmark genes. This approach enables spatial visualization of gene expression, gene set enrichment, transcriptional program activity, and cell-type annotation on tissue sections. Our framework integrates spatial information into downstream modeling and computational analysis, applicable to any tissue or dataset with single-cell or single-nucleus RNA sequencing data and limited spatial transcriptomic data. This dataset offers unprecedented resolution and depth, allowing researchers to uncover cellular and molecular processes influenced by the tissue microenvironment.

## Results

To achieve a systems-level understanding of how the stroma and parenchyma interact during kidney development, we sought to characterize the relationships among all cell types in the kidney at discrete developmental stages while considering the organ’s complex spatial architecture.

To accomplish this, we analyzed single-cell mRNA sequencing datasets from embryonic day 18.5 (E18.5) mouse kidneys.^9,16^ Using the EIGEN algorithm,^17^ we identified high-quality marker genes representing distinct cellular populations or states within the dataset (Figure S1). We selected 90 landmark genes, including multiple markers for each cell cluster, to ensure comprehensive coverage of all known cell types in the E18.5 mouse kidney. These included both previously described and novel markers. Notably, the selected genes were enriched for stromal cell markers, reflecting our specific research focus. A heatmap of the chosen landmark genes (Figure 1) illustrates distinct expression patterns across cell types, providing a robust foundation for further investigation.

**Figure 1.**
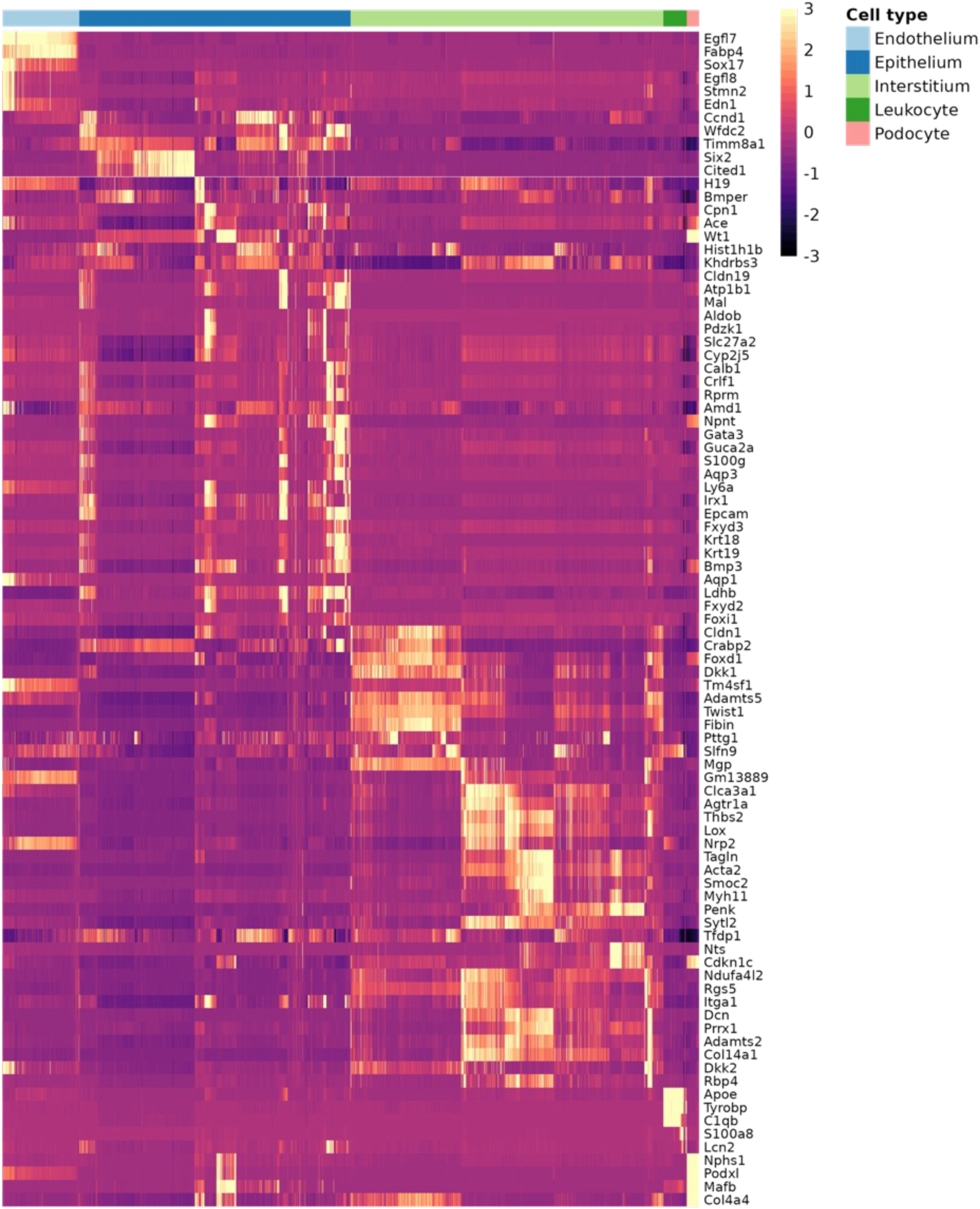
Expression of landmark genes across cell types. A heatmap showing the relative expression levels of landmark genes across different cell types. The data is from a combined single-cell sequencing dataset from e18.5 kidney (GEO accession numbers GSE108291 and GSE155794). The expression values are imputed and normalized to account for technical variability in the data. Cell types are annotated and ordered along the x-axis, as shown in the top legend.

We next performed direct RNA in situ sequencing (ISS)^18^ on multiple sections of mouse kidneys at embryonic day 15.5 (E15.5), E18.5, and postnatal day 3 (P3) using fluorescently tagged barcodes for the 90 landmark genes. This method enables direct RNA detection in tissues by combining padlock probes and rolling circle amplification with in situ sequencing chemistry. Of the 90 targeted genes, 88 produced measurable signals across all three time points. Representative data from each time point are shown in Figure S2A–C.

We analyzed single-cell and single-nucleus RNA sequencing datasets from mouse kidneys at corresponding developmental stages. Previously published single-cell RNA sequencing data for E15.5 mouse kidneys were obtained from Lawlor et al.^19^ and Naganuma et al.^20^ via the Gene Expression Omnibus (GSE118486 and GSE149134). For E18.5 and P3, we generated our own datasets, as detailed in the Methods section. Following standard preprocessing, we obtained 5,339 transcriptomes from E15.5 kidneys, 15,389 from E18.5, and 10,008 from P3. Of the original 90 landmark genes, 85, 86, and 80 genes, respectively, had sufficient spatial and single-nucleus transcriptomic data to support further analysis.

Using a statistical model that predicts the likelihood of a cell’s location based on its landmark gene expression profile, we integrated transcriptome measurements from individual nuclei with their physical locations on reference kidney sections. This approach enabled us to map transcriptional data from each cell or nucleus onto a spatial framework, effectively generating a virtual in situ hybridization image for every gene expressed in the datasets. We refer to this resource as the Kidney Spatial Transcriptome Analysis Tool (KSTAT).

To assess the accuracy of our mapping approach, we compared actual gene expression measurements from in situ sequencing (ISS) to the predictions generated by our model for the landmark genes. Since the landmark genes were selected based on clusters identified in E18.5 transcriptomic data, we focused our evaluation on this time point. The model assigns non-zero expression probabilities to multiple locations within the kidney section, resulting in diffuse calculated signals. To facilitate visual comparisons, we applied a threshold to the predicted expression data, selecting the highest-probability locations for display. The number of selected locations matched those with measured ISS signals. We observed a strong correlation between the measured and predicted gene expression data; however, some discrepancies were noted. For example, while Podxl mRNA is expressed in both podocytes and endothelial cells at E18.5, our predictions captured podocyte expression but underrepresented endothelial expression. We hypothesized that these differences arose from setting the expression threshold too high. To address this, we developed two solutions: (1) an automated method for optimizing thresholds (described in the Methods section) and (2) a user-controlled threshold adjustment feature, analogous to varying the development time in colorimetric mRNA in situ experiments. Figure 2 shows predicted expression data and the effects of applying three arbitrarily chosen thresholds (90%, 95%, and 99%) for four representative genes. Overall, thresholds in the 95–99% range most accurately reproduced the measured signals.

**Figure 2.**
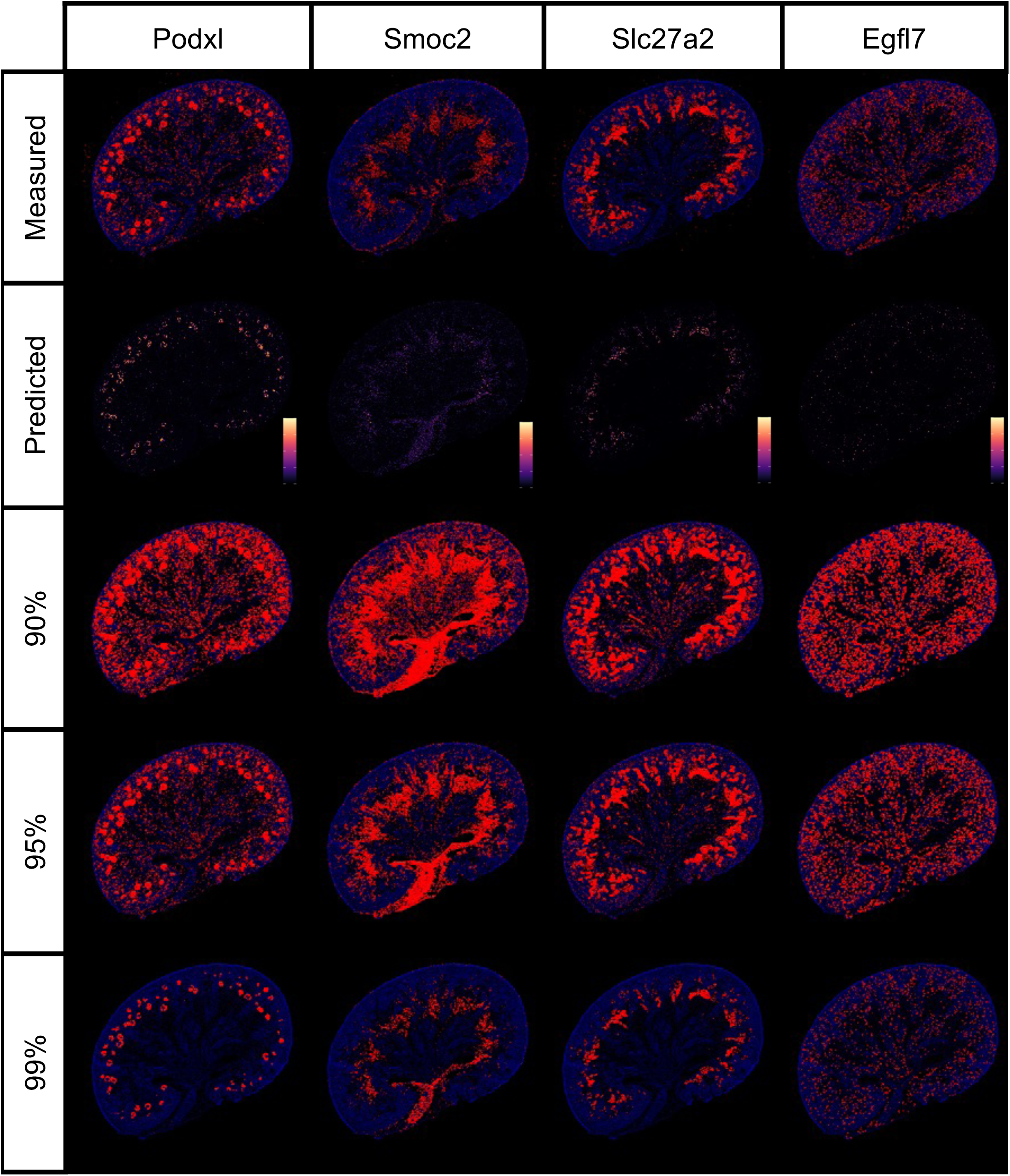
Thresholding reconstruction and thresholding. This figure compares the transcript distributions measured by CARTANA with those inferred by KSTAT for four genes: Podxl, Smoc2, Slc27a2, and Egfl7. For each gene, the expected expression calculated using KSTAT is shown alongside the results of applying thresholds at the 90th, 95th, and 99th percentiles.

We quantitatively evaluated the performance of our model by assessing the reconstruction error for each landmark gene using both the full model and leave-one-out cross-validation (LOOCV), a standard machine learning approach for estimating model generalizability to unseen data.^21^ The first analysis assessed the model’s ability to reconstruct measured gene expression using all available data, while LOOCV provided an estimate of the model’s predictive capacity for genes excluded from the training set. To quantify uncertainty and assess the fidelity of each measurement, we estimated the probability of obtaining the observed expression or a more extreme value if the signal were randomly distributed throughout the reference space (Supplemental Table 1). This was achieved by calculating the distance between the measured and inferred expression distributions and comparing these values to the distances measured for a bootstrapped random distribution. Using a lower tail probability threshold of 5%, we accurately reconstructed spatial gene expression for 86 of the 88 landmark genes. LOOCV revealed reconstruction errors for four genes, resulting in accurate spatial predictions for 84 of 88 genes, an estimated accuracy of 95% when predicting spatial distributions for genes without spatial transcriptomics data.

We further validated our approach by comparing the imputed expression of several non-landmark genes with results from standard colorimetric in situ hybridization using single-probe sections. Predicted expression patterns qualitatively matched those obtained using antisense in situ hybridization techniques (Figure 3A). Figure S3 shows predicted expression data for these genes at both E15.5 and postnatal day 3 (P3), demonstrating consistency across developmental stages.

**Figure 3.**
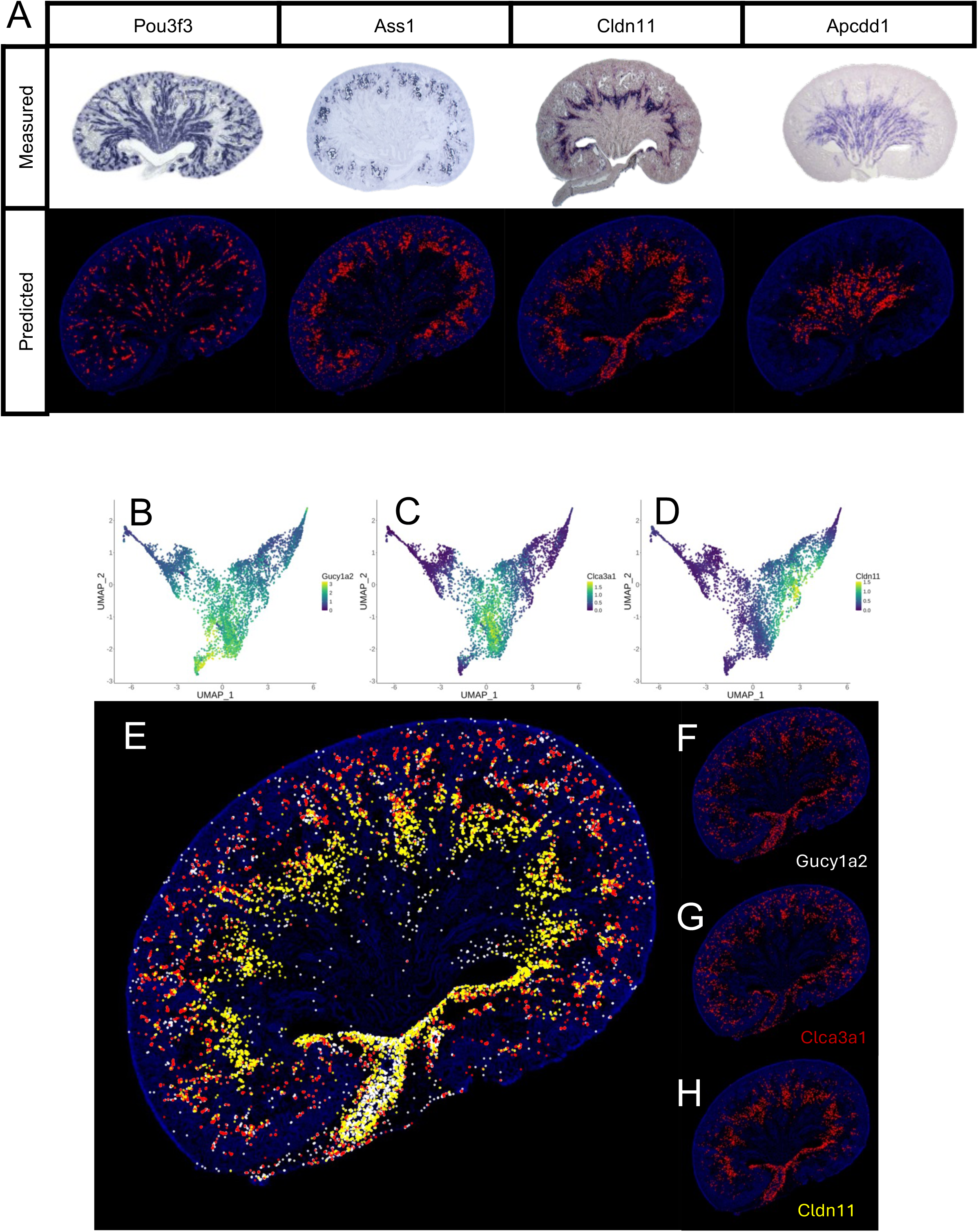
Multiplexed prediction of high-resolution spatial expression for unmeasured genes. (A) Validation of our model’s ability to predict gene expression patterns for unseen genes. The predicted spatial distributions of Pou3f3, Ass1, Cldn11 and Apcdd1 are compared to their corresponding expression patterns measured using colorimetric in situ hybridization in e18.5 kidneys. Notably, these four genes were not part of the training set, demonstrating our model’s capacity to generalize to new genes. (B-D) UMAP’s of isolated interstitial cells depicting expression of Gucy1a2, Clca3a1 and Cldn11 which are differentially enriched in three separate clusters of these cells. (E) The simultaneous projection of predicted expression for Gucy1a2, Clca3a1 and Cldn11 onto the 2D kidney expression permits appreciation of spatial relationships between multiple genes with related yet distinct expression patterns (Gucy1a2, white; Clca3a1, red; Cldn11, yellow). (F-H) Individual projections of predicted expression for Gucy1a2, Clca3a1 and Cldn11.

The raw in situ sequencing data provides subcellular resolution, with a pixel edge length of 0.16 µm, surpassing the resolution of traditional in situ hybridization methods. For visualization, the diameter of each plotted point was adjusted to enhance aesthetic clarity. Figure 3B–D shows UMAP plots for three genes—**Gucy1a2**, **Clca3a1**, and **Cldn11**—within distinct clusters of interstitial cells. Previous investigations in our lab, constrained by the resolution limits of standard colorimetric in situ hybridization, were unable to determine whether these genes exhibited overlapping expression patterns. Figure 3E simultaneously visualizes the expression of all three genes, with point diameters approximating the average mammalian cell size (15 µm) to ensure that non-overlapping points represent distinct cells. Using KSTAT, we demonstrate that **Gucy1a2**, **Clca3a1**, and **Cldn11** exhibit non-overlapping expression patterns (Figure 3F–H), underscoring the utility of our approach for resolving spatial relationships at high resolution.

In addition to mapping individual transcripts, our resource allows for the spatial mapping of transcriptional clusters. Recent studies using traditional methods have revealed transcriptional heterogeneity in endothelial cells of the embryonic kidney.^22,23^ However, similar analyses have not been conducted for vascular-associated cells, such as mural cells. Figure 4A shows a UMAP of isolated stromal cells, highlighting Pdgfrb expression, a canonical mural cell marker. Pdgfrb transcripts are enriched in four distinct transcriptional clusters. Figures 4B–E display UMAPs individually highlighting each of these clusters. The spatial relationships of these different clusters is unknown. To address this, we used KSTAT to map the clusters simultaneously (Figure 4F) and individually (Figures 4G–J) onto our E18.5 reference section. These data revealed significant spatial heterogeneity among mural cells. To validate this finding, we identified transcripts differentially enriched across the four clusters (Mgp, Cspg4, Acta2, and Abcc9). UMAPs representing expression of these genes are shown in Supplemental Figures 4A–D. Spatial mapping with KSTAT, combined with canonical mural cell markers (Rgs5 and Pdgfrb), confirmed the spatial differences in gene expression (Supplemental Figures 4E–K).

**Figure 4.**
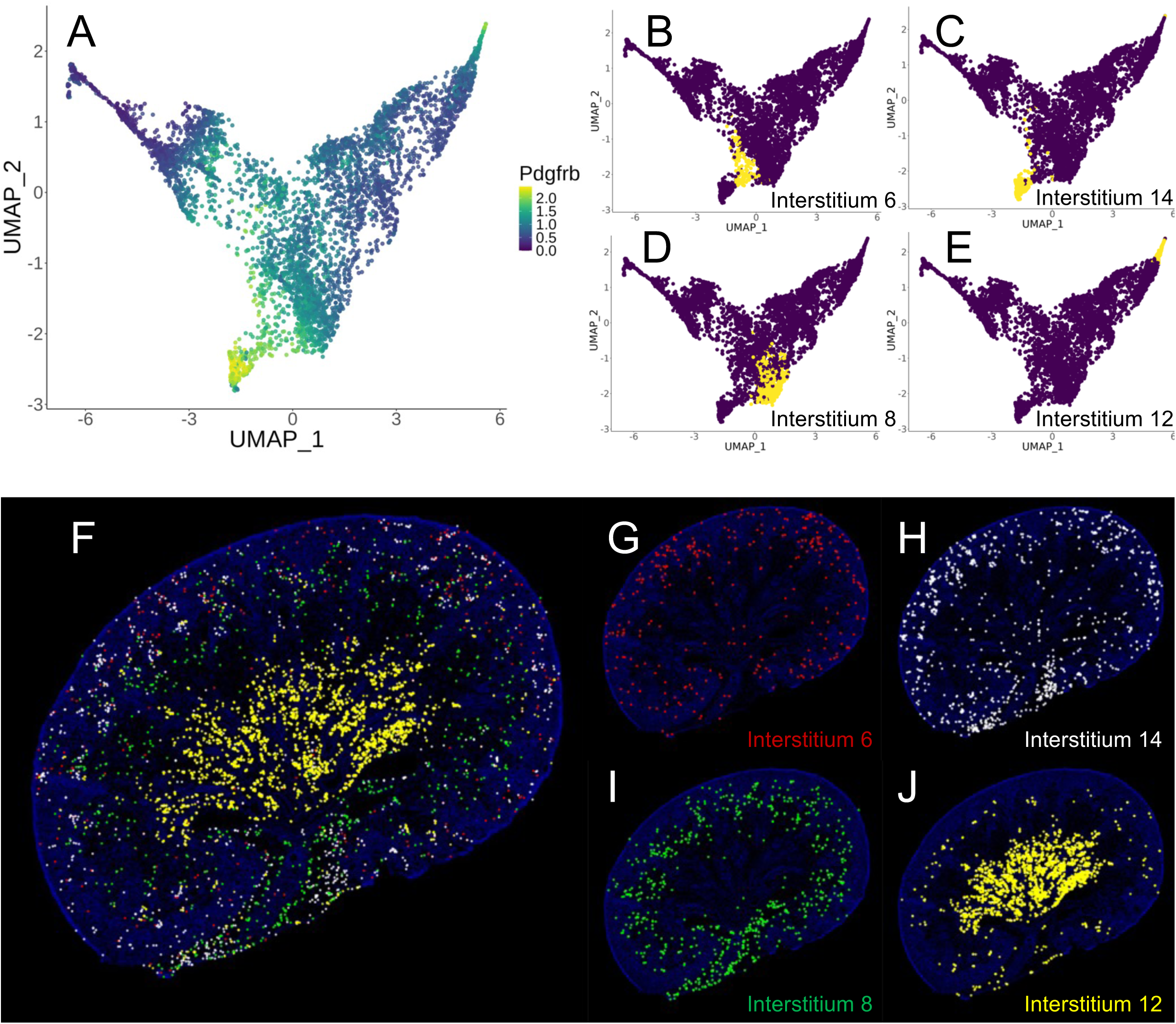
Spatial heterogeneity of mural cells. (A) UMAP of isolated stromal cells colored by expression of the mural cell marker Pdgfrb. (B-E) UMAPs indicating four snRNA-seq expression clusters predicted to be marked by Pdgfrb expression. (F) Composite prediction of the spatial locations of the four snRNA-seq expression clusters highlighted in B-E (6, red; 14, white; 8, green; 12, yellow). (G-J) Individual predictions of the predicted spatial locations of the four clusters identified in B-E.

To corroborate these findings, we performed multiplex fluorescent in situ hybridization on E18.5 kidney sections using probes for the identified clusters (Supplemental Figures 4L–P). This analysis confirmed the spatial heterogeneity of the the four clusters (Figure S4L). Additionally, the in situ data detected two known mural cell types—mesangial cells and vascular smooth muscle cells—that were not resolved as transcriptionally distinct clusters in the snRNA-seq data (see Discussion).

At E18.5, Gucy1a1 and Gucy1a2 were differentially enriched in two of the mural cell clusters discussed above. The Gucy genes encode subunits of soluble guanylate cyclase (sGC), an enzyme activated by nitric oxide that converts GTP to cGMP. This signaling system has been implicated in pericyte maintenance, proliferation, and vascular smooth muscle tone regulation.^24–26^ The sGC enzyme is a heterodimer consisting of alpha and beta subunits. In our E18.5 and P3 snRNA-seq datasets, Gucy1b1 was the only beta subunit detected and appeared to encompass the expression domains of both Gucy1a1 and Gucy1a2 at E18.5 (Figure 5A–B and Supplemental Figures 5A–C). At P3, Gucy1a1 and Gucy1a2 expression remained largely non-overlapping but became more restricted to the juxtamedullary and outer medullary regions, respectively (Figure 5C–D and Supplemental Figures 5D–F). Gucy1b2 expression overlapped almost entirely with Gucy1a2 and showed minimal overlap with Gucy1a1. These observations suggest that distinct isoforms of the alpha and beta subunits are under separate transcriptional control and may contribute to the development/function of specific pericyte populations and/or distinct developmental processes.

**Figure 5.**
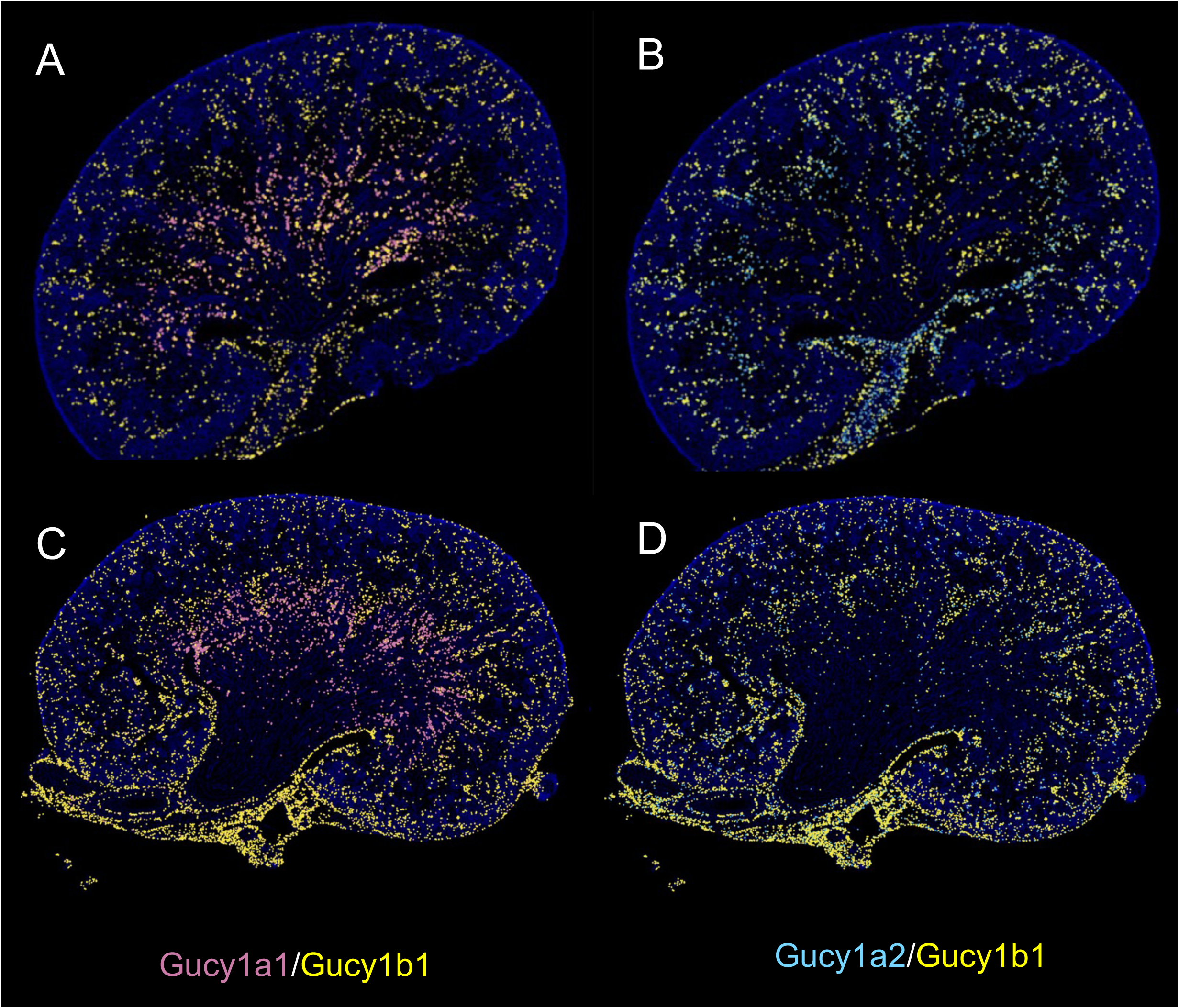
Late embryonic and early postnatal heterogeneity of soluble guanyl cyclase expression.

Our resource can also assign spatial information to predicted transcription factor activity (regulon activity). For example, using SCENIC^27^, we predicted Lef1 regulon activity from snRNA-seq data and projected it onto the E18.5 UMAP (Figure 6A). Spatial mapping of the Lef1 regulon onto the reference tissue section (Figure 6B) revealed predicted activity in the medullary stroma, cortical renal vesicles, and ureteric bud.

**Figure 6.**
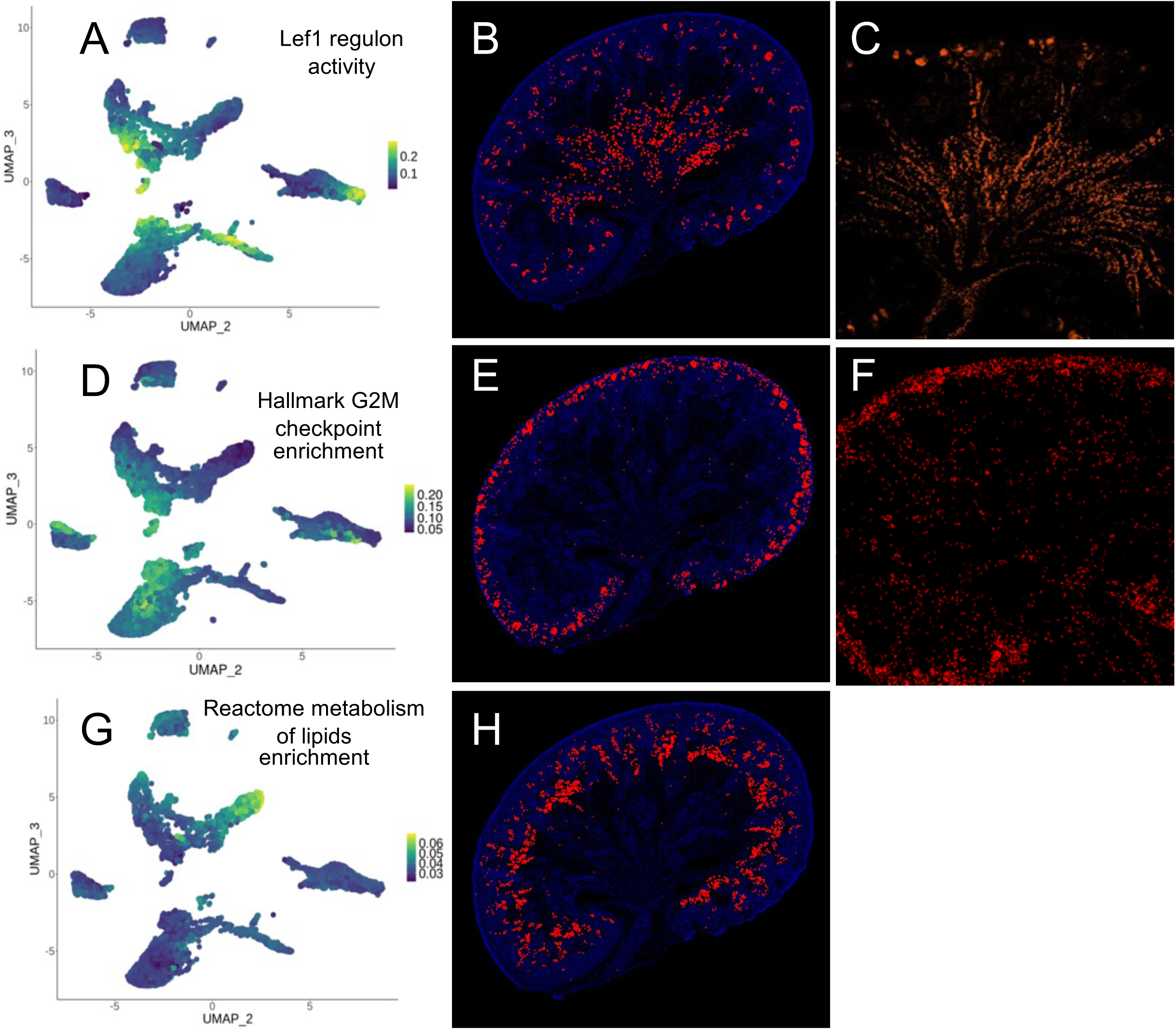
Projection of cell properties. (A) UMAP depicting the results of regulatory network analysis using SCENIC reveals Lef1 regulon activity (B) The corresponding spatial distribution of highest predicted Lef1 regulon inferred by KSTAT. (C) Inmmunoflourescence localization of Lef1 in an E18.5 kidney section. (D) Enrichment analysis of MSigDB hallmark genes involved in the G2/M checkpoint projected onto a UMAP. (E) The predicted spatial enrichment of this gene with a threshold applied to show the top percentile. (F) EdU labeling after a 1.5-hour pulse in E18.5 mouse kidney (EdU, red). (G and H) Enrichment analysis of lipid metabolism-related genes measured with AUCell (left, UMAP plot). The predicted spatial enrichment of this gene set is shown on the right, with a threshold applied to display the top percentile.

This spatial pattern closely mirrors both the expression of Lef1 (Figure 6C) and previously demonstrated activity of LEF1 protein in the kidney^28–30^.

The KSTAT resource also enables cell-level mapping of gene set enrichment for curated gene sets, such as those available in MSigDB^31^. For instance, in Figure 6D, we mapped the hallmark gene set for G2 to M progression onto the E18.5 UMAP. Figure 6E projects the gene set enrichment onto the reference kidney section. Enrichment of the G2 to M gene set in the kidney cortex aligns with known indicators of cell division. This finding was corroborated by EdU incorporation assays, where a two-hour EdU pulse administered to pregnant dams revealed active cell division in the cortical region (Figure 6F).

Next, we evaluated whether metabolic pathways could be spatially mapped in the developing kidney using transcriptomic data. In Figure 6G, we mapped the Reactome “metabolism of lipids” gene set onto the E18.5 UMAP. Using AUCell^27^ to measure gene set enrichment in individual nuclei, we projected the results onto the reference tissue section (Figure 6F). The data suggest that lipid metabolism is active in maturing proximal tubules, consistent with recently published findings.^32^

The early stages of kidney development rely on interactions between mesenchymal and epithelial cells. Our dataset enables the investigation of cell-cell communication by modeling ligand-receptor interactions while incorporating spatial information within the tissue architecture. To test the utility of our dataset in identifying biologically significant interactions, we queried ligand-receptor pairs predicted to be active within the metanephric mesenchyme (MM). Initially, we used CellPhoneDB, an algorithm that does not account for spatial information.^33^ Among the predicted interactions was the ligand Wnt5a binding to the co-receptor complex Fzd4/Lrp6 (Table S2). Our E18.5 spatial atlas revealed that Wnt5a is expressed in the medullary stroma, while Fzd4/Lrp6 is co-expressed in cortical nephron progenitor cells (Figure 7A), consistent with previous findings.^29^

**Figure 7.**
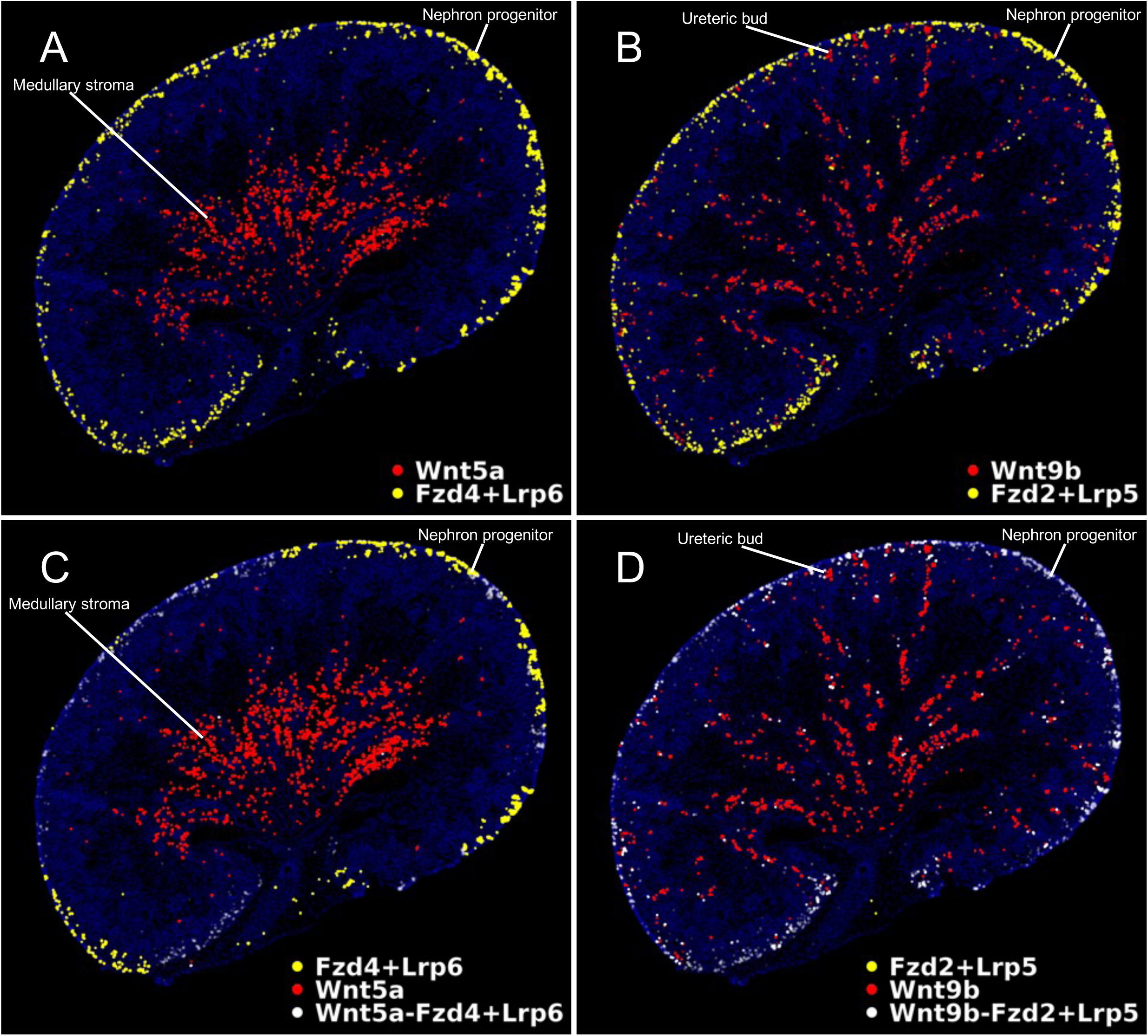
Predicting ligand-receptor interactions with KSTAT. This figure demonstrates the ability of our model (KSTAT) to predict ligand-receptor interactions in the kidney. (A and B) The predicted spatial distributions of ligand-receptor complexes inferred by KSTAT are shown for two interactions: Wnt5a with co-receptors Fzd4 and Lrp6, and Wnt9b with Fzd2 and Lrp5. (C and D) The results of optimally transporting the ligands to their receptors are displayed. Locations where receptor complexes receive ligand are highlighted in white, with opacity determined by the ratio of ligand to receptor concentration. This visualization illustrates the predicted spatial patterns of ligand-receptor binding.

We next used our spatial dataset to refine the analysis by incorporating spatial expression patterns of the interacting proteins. Interestingly, our model deprioritized the Wnt5a/Fzd4 interaction and instead predicted that Wnt9b signals through Fzd2/Lrp5. The spatial dataset confirmed that Wnt9b is expressed throughout the ureteric bud and collecting ducts, while Fzd2/Lrp5 is co-expressed in nephron progenitors (Figure 7B). Previous genetic work suggests that Wnt9b, and not Wnt5a, signals to nephron progenitors.^6,7,34^ Nonetheless, we used the data from KSTAT to quantitatively assess the likelihood of these interactions. Weutilized GeomLoss^35^ to calculate the spatial distances between cells producing the ligands and receptors, reasoning that the likelihood of interaction is inversely related to this distance.

Figures 7C and 7D depict the predicted receptor-bound ligands, with bound receptors appearing white and unbound receptors appearing yellow. Saturation levels are represented by the opacity of the spots denoting bound receptors: low predicted ligand binding is nearly transparent, while stoichiometric saturation is shown as bright white. Figure 7C illustrates minimal effective binding for the Wnt5a/Fzd4-Lrp6 complex, whereas the Fzd2/Lrp5 receptor complex is almost fully saturated with Wnt9b (Figure 7D). This analysis is consistent with previously published work and underscores the importance of spatial information in improving predictions of biologically relevant cell-cell communication events.

## Discussion

Recent advances in transcriptomic technologies have enabled the creation of gene expression atlases for various tissues and organs. Traditional approaches, such as antisense mRNA in situ hybridization and antibody staining, provide spatial context but are limited by shallow gene coverage and variability between tissue sections, complicating comparisons across genes. Conversely, single-cell transcriptomic datasets offer greater depth, capturing thousands of sequencing reads per cell and profiling thousands of cells simultaneously. However, tissue dissociation in these methods results in a complete loss of spatial context, hindering analyses that rely on understanding cellular relationships within the native tissue architecture. Spatial transcriptomics bridges this gap, enabling simultaneous measurement of multiple transcripts directly in tissue. However, existing platforms are constrained by trade-offs between spatial resolution and transcriptome coverage. Our study addresses these challenges by integrating single-cell transcriptomic data with high-resolution spatial mapping, creating a comprehensive atlas of gene expression in embryonic and postnatal mouse kidneys. This integrative approach provides novel insights into cellular heterogeneity, spatial gene expression, and cell-cell communication in the kidney.

To overcome the limitations of existing technologies, we developed the Kidney Spatial Transcriptome Analysis Tool (KSTAT), which combines the depth of single-cell/nucleus sequencing data with spatial transcriptomics. Using a targeted set of cell-type-restricted landmark genes, KSTAT enables spatial assignment of transcriptomic data to (sub)cellular locations within reference E15.5, E18.5, and P3 kidney sections. This integration generates a near-comprehensive atlas of gene expression while also facilitating the correlation of spatial location with advanced analyses such as cellular annotations, gene set enrichment, and regulon activity. Leveraging KSTAT, we explored mural cell heterogeneity, a feature critical for understanding disease states and advancing tissue engineering efforts for renal replacement therapies. Our findings revealed previously unrecognized heterogeneity among kidney mural cells at E18.5. Notably, validation of our predictions suggests that the true diversity of these cells may exceed our initial computational predictions.

Current methods for visualizing single-cell ‘omics data typically rely on dimensionality reduction techniques like tSNE or UMAP, which position datapoints (cells) on a two-dimensional map. While distances between datapoints can indicate relationships between cells, these plots provide no information about the spatial location of cells within the tissue. This low-dimensional representation is useful for some analyses but lacks the critical spatial context needed for others, such as receptor-ligand interaction studies. Without spatial data, users must rely on prior knowledge or annotations to infer relationships between cells, a process that is both labor-intensive and prone to error. KSTAT overcomes this limitation by integrating spatial information, enabling automated, unbiased mapping of transcriptomic data to tissue architecture. This is particularly valuable for researchers new to a model system and for studying pathological tissues, where novel cell states may emerge. By incorporating spatial data into the analysis pipeline, KSTAT expands the utility of single-cell transcriptomics, providing a robust framework for interrogating cellular behavior in both health and disease.

Our validation demonstrates that integrating single-cell or single-nucleus RNA sequencing with limited spatial transcriptomics can accurately predict gene and pathway expression at a near-comprehensive level. However, there are important considerations for researchers aiming to generate similar resources. Chief among these is achieving a high success rate with in situ sequencing, which depends on several factors. First, careful selection of landmark genes is critical. Landmark genes must generate meaningful data, meaning their expression should be detectable but not ubiquitous. Broadly expressed genes must not saturate detection. To address this, we used our previously published algorithm to select landmark genes that fall within this “sweet spot” of detectability ^17^.

In our experiment, we successfully detected 88 of the 90 targeted genes, achieving a high success rate. The two undetected genes were likely excluded due to either low expression levels or technical issues during sequencing or visualization. Among the detected genes, four (H19, Fxyd2, Acta2, and Ldhb) exhibited broad expression patterns. This likely reflects enrichment in specific cell types combined with lower-level expression in others. Re-examination of our single-nucleus RNA-seq data revealed that these genes had relatively high proportions of cells expressing them above a certain threshold relative to intercluster variance. Interestingly, genes with more extreme deviations (S100g, Fabp4, Wfdc2, and Mgp) displayed more spatially restricted expression in the in situ sequencing data, suggesting potential limitations in using single-nucleus RNA-seq to predict saturation in spatial transcriptomics. Despite this, we obtained spatially restricted expression for 86 of the 90 genes, and the saturated signal from these four genes did not significantly impact results due to the redundancy of markers for each cell type. For instance, the imputed expression of Acta2 closely matched its known expression pattern.

The success of this technique depends not only on high-quality in situ sequencing but also on high-quality transcriptomic data. The greater the depth and quality of the single-cell transcriptomic data, the more comprehensive the resulting atlas will be. Genes or cell types absent from the single-cell dataset cannot be mapped. For instance, the E18.5 kidney dataset used to select landmark genes was enriched for stromal cells but lacked sufficient representation of ureteric and endothelial cells, limiting its capacity to capture pathways active in these compartments. Similarly, since landmark genes were derived from E18.5 data, their representation at E15.5 or P3 may be less robust. Additionally, our single-nucleus sequencing data was generated from whole kidneys without enrichment for specific cell types, potentially underrepresenting rare populations. This may explain the absence of distinct clusters for mesangial cells or vascular smooth muscle cells, which were observed through multiplex in situ hybridization. Mesangial cells, in particular, are challenging to dissociate from intact glomeruli, complicating their capture.

Alternatively, reparameterization of the clustering algorithm could reveal these clusters. Ideally, an exhaustive single-cell dataset with at least 5–6 landmark genes for every cluster across all time points would maximize accuracy. However, such an approach must balance cost with analytical depth. While other spatial transcriptomics platforms, such as the 10x Visium platform, offer lower resolution that may hinder certain analyses, our use of the Cartana platform—now incorporated into 10x Xenium technology—has provided high-resolution data suitable for this work.

Spatial information is particularly critical for inferring intercellular communication. Interactions involving membrane-bound ligands and receptors rely on close cellular apposition, while the range of secreted ligand activity depends on the biochemical properties of the extracellular environment. Although computational modeling of signaling relationships based on molecular expression has advanced, incorporating spatial data into such analyses remains a nascent field. Existing spatial transcriptomics platforms often impose limitations on data depth and resolution, constraining their utility. By integrating single-cell and spatial transcriptomics data with computational optimal transport methods, KSTAT overcomes these obstacles. For example, mapping probable Wnt-pathway ligand and receptor locations in the developing kidney demonstrates how spatial data can refine hypotheses about signaling interactions. This integration represents a significant step forward in understanding cell-cell communication, offering insights into potential therapeutic targets and strategies.

By integrating spatial data with single-cell transcriptomics, KSTAT enables researchers to generate specific hypotheses about tissue development. However, beyond kidney development, this study provides a roadmap for how to generate additional expression atlases. In pathological contexts, novel cell states often emerge and influence disease progression. Our approach allows precise mapping of disease-associated transcriptional programs or cell states within tissues, providing a foundation for identifying spatially localized signaling pathways. This knowledge can inform the development of targeted therapeutic strategies by pinpointing communication networks that regulate key transcriptional programs. Ultimately, these types of resources will open up new avenues for understanding and modulating development, maintenance, regeneration and disease, offering powerful tools for advancing precision medicine.

## Resource Availability

### Lead contact

Further information and requests for resources and reagents should be directed to and will be fulfilled by the lead contact, Thomas Carroll.

### Materials availability

This study did not generate new unique reagents.

### Data and code availability

- Single-nucleus RNA-seq data have been deposited at GEO and are publicly available as of the date of publication.
- All original code has been deposited at Zenodo and is publicly available as of the date of publication. DOIs are listed in the key resources table.
- Any additional information required to reanalyze the data reported in this paper is available from the lead contact upon request.

## Acknowledgements

We would like to thank Dr. Morgane Rouault, Dr. Courtney Karner, Dr. Robert Tower and members of the Carroll, Cleaver and Marciano laboratories for reading and providing comments on this study and manuscript. Financial support for this work provided by the NIH/NIDDK, grants RC2DK125960 and R01DK127634, and the NIDDK Diabetic Complications Consortium (DiaComp, www.diacomp.org), grant DK076169.

## Author Contributions

Conceptualization: Christopher P. Chaney, Thomas J. Carroll

Data curation: Christopher P. Chaney, Alexandria N. Fusco, Elyse D. Grilli, Jane N. Warshaw, Peter M. Luo

Formal analysis: Christopher P. Chaney

Funding acquisition: Ondine Cleaver, Denise K. Marciano, Thomas J. Carroll

Investigation: Christopher P. Chaney, Alexandria N. Fusco, Elyse D. Grilli, Jane N. Warshaw, Peter M. Luo

Methodology: Christopher P. Chaney, Thomas J. Carroll

Project administration: Thomas J. Carroll

Software: Christopher P. Chaney

Resources: Ondine Cleaver, Denise K. Marciano, Thomas J. Carroll

Supervision: Ondine Cleaver, Denise K. Marciano, Thomas J. Carroll

Validation: Christopher P. Chaney, Thomas J. Carroll

Visualization: Christopher P. Chaney, Thomas J. Carroll

Writing – original draft: Christopher P. Chaney

Writing – review & editing: Christopher P. Chaney, Thomas J. Carroll

## Declaration of interests

The authors declare no competing interests.

## Declaration of generative AI and AI-assisted technologies

During the preparation of this work the author(s) used llama3 in order to improve readability and language of the manuscript. After using this tool/service, the author(s) reviewed and edited the content as needed and take(s) full responsibility for the content of the published article.

## Supplemental information titles and legends

Document S1. Figures S1-S11 and Tables S1-S2.

## STAR Methods

### Nuclei Isolation and sequencing

All animals were housed, maintained, and used according to National Institutes of Health (NIH) and Institutional Animal Care and Use Committees (IACUC) approved protocols at the University of Texas Southwestern Medical Center (OLAW Assurance Number D16-00296). To isolate nuclei from E18.5 kidneys, we modified a protocol from Ben Humphrey’s lab.^36^ Kidneys were dissected in cold PBS without calcium or magnesium, snap-frozen for genotyping, and then thawed on ice for pooling. We minced the kidneys using razorblades and homogenized them with a Dounce homogenizer in Nuclei EZ Lysis buffer (Sigma) supplemented with protease inhibitor (Roche), RNase inhibitor (Promega and Life Technologies). The homogenate was filtered through a 40μm cell strainer (pluriSelect) and centrifuged at 500 x g for 5 min at 4°C. The pellet was resuspended, washed, and filtered again through a 5μm cell strainer (pluriSelect).

The resulting nuclei suspensions were submitted to UTSW Next Gen Sequencing Core for library preparation and sequencing using the Chromium Single Cell 3’ Prime Gene Expression kit (10x Genomics) on an Illumina NextSeq 2000.

### Single-cell (nucleus) RNA-seq preprocessing

Cells (nuclei) were called from empty droplets by testing for deviation of the expression profile for each cell from the ambient RNA pool.^37^ Cells (nuclei) with low numbers of detected genes or library sizes, as defined as a deviation greater than three median absolute deviations (MADs) below the median, were excluded from further analysis. Similarly, cells (nuclei) with large mitochondrial proportions, i.e., more than 3 mean-absolute deviations above the median, were removed. However, in the case of stripped nuclei, we enforced a minimum difference between the threshold 0f 0.5% between the threshold for exclusion and the median mitochondrial transcript count. Cells (nuclei) were pre-clustered; a deconvolution method was applied to compute size factors for all cells^38^ and normalized log-expression values were calculated.

### Landmark selection

Cells were clustered by building a shared nearest neighbor graph, which represents cells as nodes connected by edges based on their similarity.^39^ The Walktrap algorithm^40^ was then applied to identify clusters of cells with similar characteristics. After coarsely annotating the cells, stromal cells were further sub-clustered. In total, 52 clusters were found, 17 of which were stroma. To identify genes that specifically mark populations of sequenced cells, we employed the EIGEN algorithm, which leverages the “wisdom of the crowds” principle. This involves measuring pairwise differential expression between each pair of clusters using multiple statistical methods (pairwiseTTest, pairwiseBinom, and pairwiseWilcox from the scran Bioconductor package, as well as the zlm method from the MAST Bioconductor package). A manual curation step was performed to ensure both specific expression and coverage of all cell states represented in the sequencing data.

### In situ sequencing

Tissue sections were collected and pre-processed according to established protocols provided by CARTANA, a platform specializing in direct RNA targeted in situ sequencing. CARTANA has since been acquired by 10x Genomics. Of the genes selected for analysis, data could not be acquired for Amd1 and Hist1h1b due to technical limitations. In addition to the selected genes, the vendor additionally provided data for the “housekeeping” gene H1f5.

### Preprocessing

Reads were provided by CARTANA as fractional coordinates of segmented spots in a 16,619 x 14,810- pixel reference space, which represents the spatial distribution of gene expression within the tissue section. The data was processed as a binary array, where each element (x, y, l) indicates whether landmark gene l is expressed at location (x, y). To reduce computational complexity and sparsity, we aggregated data from 16 x 16-pixel tiles, effectively reducing the height and width to 926 and 1,039 bins, respectively. This step allowed us to identify 32,157 unique bit vectors of landmark expression, with different combinations occurring at varying frequencies.

### Probability inference

We analyzed snRNA-seq from E18.5 kidney that was collected separately from the scRNA-seq data used for landmark identification as above. A UMAP of this dataset annotated by cell type is shown in Figure S7. Two of the landmark genes had no reads in the snRNA-seq data, which led to their exclusion from further analysis due to insufficient information.

Following Satija et al.,^41^ we modeled the expression of each landmark gene in the snRNA-seq dataset as a mixture of normal distributions. However, our approach differed from theirs in terms of data imputation and mixture component inference. We began by representing the marginal distribution of expression for each landmark gene in the snRNA-seq dataset as a two-component mixture of normal distributions.

Droplet capture sequencing results are represented by integer counts, demonstrate sparsity and have been observed to be zero-inflated, all of which are barriers to analysis with Gaussian models. These issues are partially addressed during normalization. We observed that imputation on normalized data using Markov Affinity-based Graph Imputation of Cells (MAGIC)^42^ smoothed the data and made it much more amenable to assumption of normality, see Figure S8.

For each of the 88 landmark genes analyzed, we fit Gaussian mixtures with k components for k in {2, 3,…, 8}. The component with the highest mean was chosen to represent cells that expressed the landmark gene, the “on” population, and the remaining components were combined to represent cells that did not express the gene, the “off” population. At the completion of this stage, we possessed estimates of the mean and covariance in “on” and “off” populations for each landmark gene, c.f. Figure S9A.

Next, we estimated multivariate normal distributions representing the probability of a cell occupying each bin as a function of that cell’s expression of the landmark genes and the measured landmark signature for that bin. For each landmark, we chose the parameters for its marginal distribution from those inferred above according to whether there was signal from the landmark measured at that location. This readily yielded an estimate of the l-dimensional mean vector. The covariance was approximated by a matrix with the variances calculated for each landmark’s selected component along the diagonal.

With the inferred distributions corresponding to each bin with a unique landmark signature in hand, calculation of the likelihood that a cell was located at a specific bin was reduced to an embarrassingly parallel computation of multivariate density. The likelihood of each transcriptome occupying the bin was then calculated using the *dmvnorm* function from the *mixtools*^43^ R package.

Ultimately, one obtains a (*N* x *B*)-dimensional tensor encoding an estimate of the likelihood of each of the *N* cells being found at each of the B unique bins in the *H* x *W* reference space. To permit further computation normalization to yield proper probability distributions for each unique bin was required.

Rather than application of *softmax* or sum-normalization which would result in full support over the population of cells, we employed *sparsegen-lin,*^44^ an adaptation of the *sparsemax* transformation,^45^ that allows control of the degree of sparsity. The inferred probabilities were broadcast to all bins with the same landmark expression signature yielding an *N^c^* x *N^b^* dimensional matrix *P* where *N^c^* is the number of transcriptomes (cells or nuclei) and *N^b^* is the number of bins.

### Projection of cell features

With the above inferred probability distributions in hand, we can now proceed to project various properties of the cells onto the reference space. This involves calculating an expectation over each bin’s distribution, which enables us to quantify specific features of interest, see Figure S9B. For instance, the expected expression of gene g at pixel i is given by

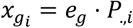

Since the probability distribution is inferred bin-wise, transitions between adjacent pixels are inherently abrupt. This leads to high-frequency fluctuations in the signal, which can be problematic. To address this, we reduced noise by applying a Gaussian filter to the expectation suppressing high frequencies while simultaneously minimizing spatial spread. The signal could then be visualized as the total expectation (Figure 2, second row) or a threshold applied to yield a virtual *in situ* hybridization image as depicted for three different thresholds in the lower part of Figure 2.

In addition to projecting continuous features, we can also project categorical properties of cells using a similar approach. This involves encoding whether a given cell belongs to a particular category, which can be represented as a binary-valued indicator function supported by the cells. As an example, transfer of the annotation of cells in the sequencing data set as those forming the proximal tubule are encoded as a binary vector with one indicating the presence of the annotation and 0 otherwise. On right-multiplying by the cells x bins probability matrix, one obtains the expected value of the indicator function at each pixel which is proportional to the likelihood that an annotated cell corresponds to that location. This calculated expression can be used for further downstream analysis or binarized by thresholding for visualization (Figure S9C).

### Automatic thresholding

We developed an automatic thresholding procedure based on the observation that the inferred signal, when ordered in ascending order, typically assumes an inverted sigmoid shape. To identify the elbow point of this curve, we implemented a method that requires a user-defined hyperparameter k. First, we sorted the unique non-zero expected values in ascending order. Then, we fit two linear models: one to the k smallest values starting from the median point, and another to the k largest values. The intersection point of these two lines was calculated, and the corresponding expected value at this point was identified as the threshold. The results of applying this thresholding procedure to the landmark Nphs1 are illustrated in Figure S10.

### Assignment of probability of marking by Pdgfrb

To determine that a cluster was marked by Pdgfrb expression, we conducted the following analysis. The data included the imputed expression of Pdgfrb all of the N cellsand the assignment of each cell to one of J clusters. We assumed the mean expression of Pdgfrb in each cluster, uk, to be normally distributed, the unknown threshold that separated clusters marked by expression of Pdgfrb from those that do not, T, was normally distributed and the slope controlling the sharpness of the decision boundary, alpha, was Cauchy distributed.

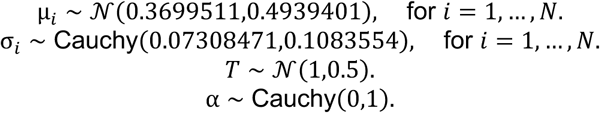

The mean and standard deviation to parameterize the prior distribution of cluster means and the median and median absolute deviation to parameterize the prior distribution of cluster standard deviations were estimated from the expression of all genes in all cells. Non-informative priors were used for T and alpha as there was no information with which to refine the assignments.

The likelihood of the expression of Pdgfb in cell j was modeled as

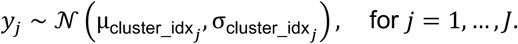

Finally, the making probability for each cluster was defined as

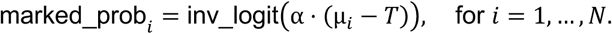

Where

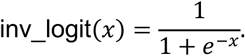

The posterior distribution of the marking probability for each cluster was sampled 10,000 times and the estimates of the mean and 89% credible intervals are displayed in Figure S11.

### Performance evaluation

To quantitatively assess the performance of KSTAT, we measured the reconstruction error between the measured landmark distribution of signal and that inferred using KSTAT. To reduce computational complexity and normalize the comparisons, once we inferred expression for a given landmark gene, we calculated a threshold so that the binarized signal would match the cardinality of the measured landmark signal. We used the Sinkhorn divergence to quantify this distance between the distributions. While this allowed us to quantify the ability of KSTAT to faithfully reconstruct the landmark signal with the fully trained model, it did not inform the likelihood of successful inference on data not previously seen by KSTAT. Thus, we also performed LOOCV. That is for each landmark, we inferred a model from all remaining landmarks and used this model to infer the left-out landmark’s distribution and then measured the error between the inferred and measured values as above. We approximated the probability of obtaining the distribution inferred by KSTAT with a bootstrap distribution derived separately for each landmark. In each case 10,000 bins were sampled from a uniform distribution supported by the two-dimensional reference space and the Sinkhorn divergence between the sampled and measured distribution was calculated. The size of the sample for each landmark was chosen to match the cardinality of the landmark signal. The lower tail mass of this distribution determined by either the reconstruction or LOOCV divergences provided an estimate of the likelihood of observing the divergence by chance. These quantitative evaluation methods provided a comprehensive assessment of KSTAT’s performance and robustness.

### Cell-cell communication inference

We investigated cell-cell communication from the perspective of receiving cells, focusing on the distribution of each component of curated receptor complexes in the population of these cells. Our analysis was restricted to ligand-receptor interactions curated in the CellPhoneDB v5.0 database. To ensure accurate calculations, we imposed stoichiometric constraints by taking the minimum expression across receptor components for each cell.

Next, we integrated the abundances of all ligand and receptor components with the debiased Sinkhorn divergence between the ligand and receptor distributions, which serves as a low-cost approximation to the earth mover’s (Wasserstein) distance. To model the spatial constraints of ligand diffusion, we used the reach parameter of the geomloss Sinkhorn SampleLoss instance to account for the limited ability of a ligand species to diffuse through space.

Our approach assumes that the cellular source of ligand does not influence whether an interaction occurs; instead, receiving cells integrate signals from all sources within their receptive field. This perspective allows us to focus on the transport of ligand species between locations in the reference space, without making assumptions about a cell’s capacity for interaction. As such, our method provides a foundation for more sophisticated models that can incorporate additional factors, such as species competition and kinetics of ligand-receptor binding. However, we only considered the ligand mass located within the distance of receptor signal specified by the reach parameter.

Key Resources Table

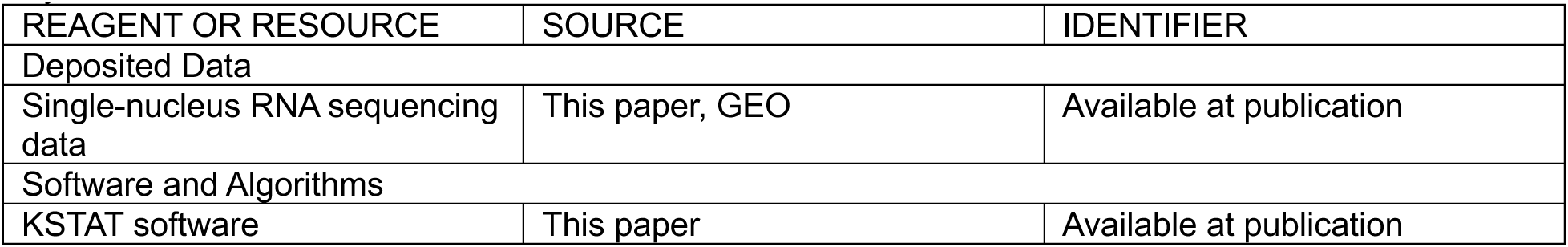

| REAGENT OR RESOURCE | SOURCE | IDENTIFIER |
| --- | --- | --- |
| Deposited Data |  |  |
| Single-nucleus RNA sequencing data | This paper, GEO | Available at publication |
| Software and Algorithms |  |  |
| KSTAT software | This paper | Available at publication |

## Notes

### Competing Interest Statement

The authors have declared no competing interest.

