## Supplemental Figures and Tables for "Integration of spatial and single nucleus transcriptomics to map gene expression in the developing mouse kidney"

### Figure S1

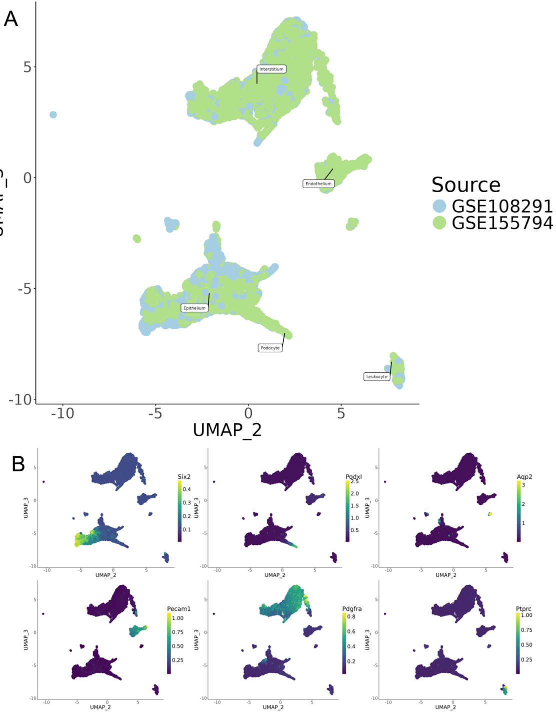
A
Interstitium
Endothelium
Epithelium
Podocyte
Leukocyte
UMAP_3
UMAP_2
Source
GSE108291
GSE155794
B
Six2
Podxl
Aqp2
Pecam1
Pdgfra
Ptprc

### Figure S2A

E15.5

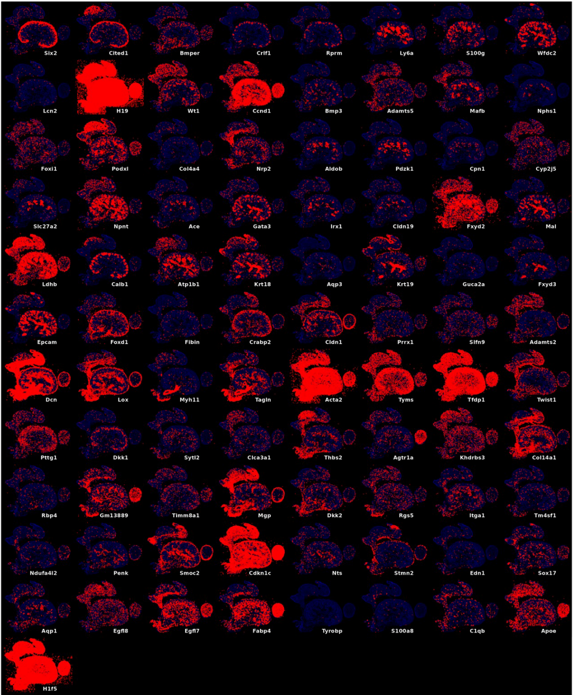

### Figure S2B

E18.5

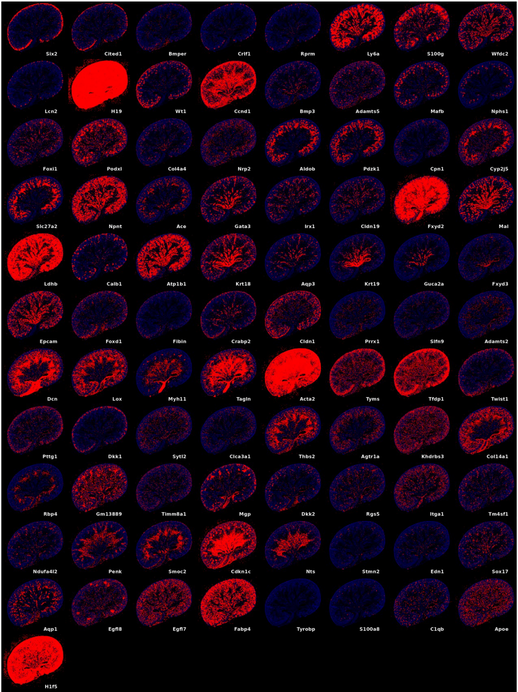

### Figure S2C

P3

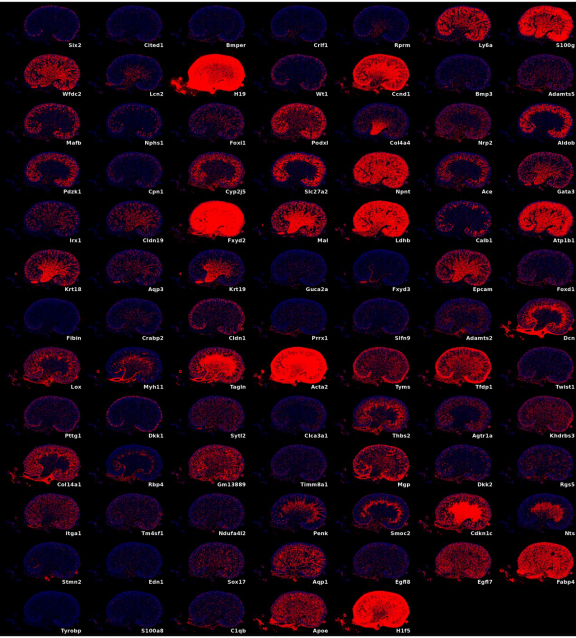

### Figure S3

E15.5

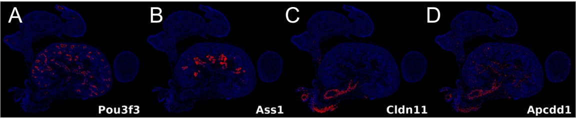


P3

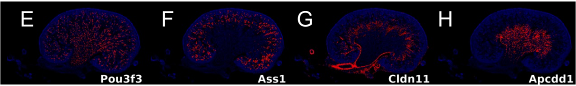

### Figure S4

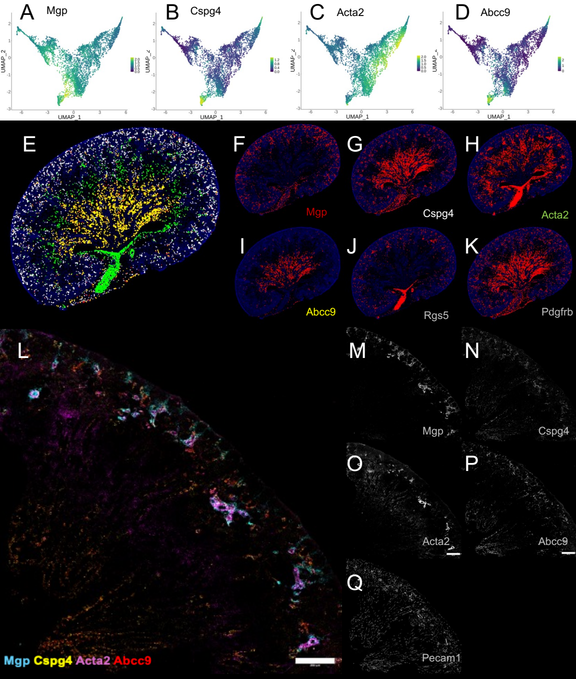
A
Mgp
B
Cspg4
C
Acta2
D
Abcc9
UMAP_1
UMAP_2
E
F
Mgp
G
Cspg4
H
Acta2
I
Abcc9
J
Rgs5
K
Pdgfrb
L
Mgp Cspg4 Acta2 Abcc9
M
Mgp
N
Cspg4
O
Acta2
P
Abcc9
Q
Pecam1

Figure S5

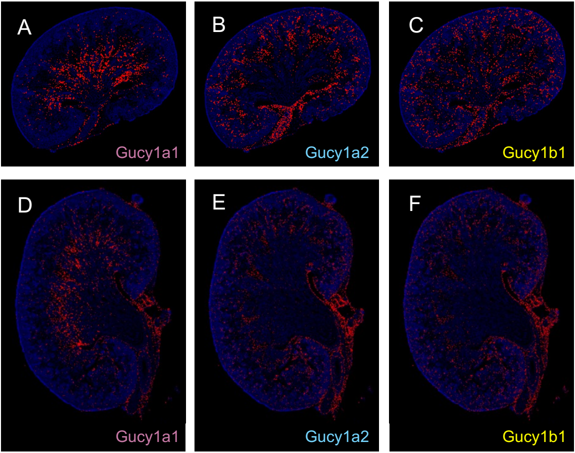

### Figure S6

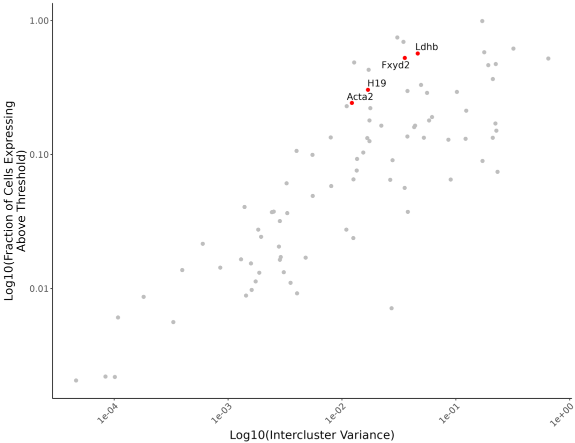

1.00
0.10
0.01
Log10(Fraction of Cells Expressing Above Threshold)
Ldhb
Fxyd2
H19
Acta2
1e-04
1e-03
1e-02
1e-01
1e+00
Log10(Intercluster Variance)

### Figure S7

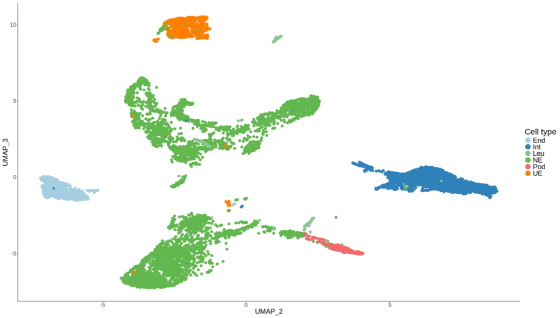

### Figure S8

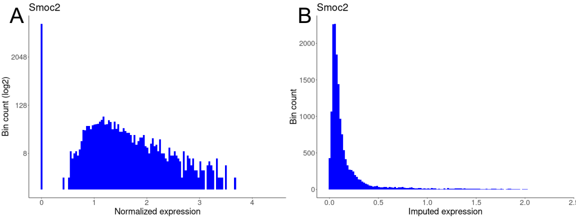
A
Smoc2
Bin count (log2)
2048
128
8
0
1
2
3
4
Normalized expression
B
Smoc2
Bin count
2000
1500
1000
500
0
0.0
0.5
1.0
1.5
2.0
Imputed expression

### Figure S9

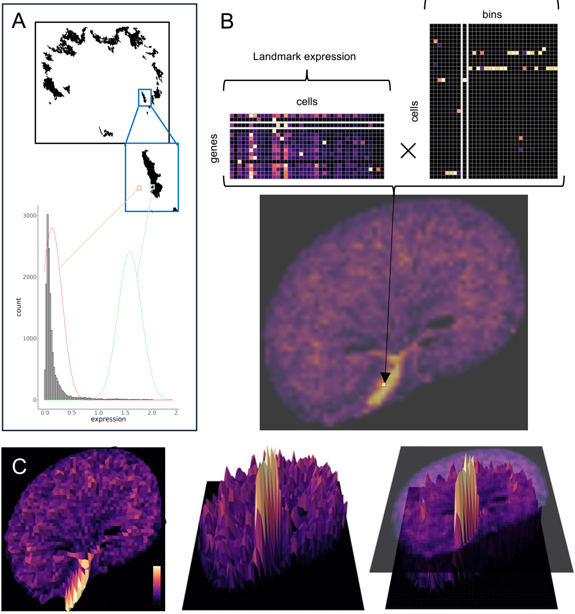
A
3000
2000
1000
0
count
0.0
0.5
1.0
1.5
2.0
2.
expression
B
Probability
bins
Landmark expression
cells
genes
cells
C

### Figure S10

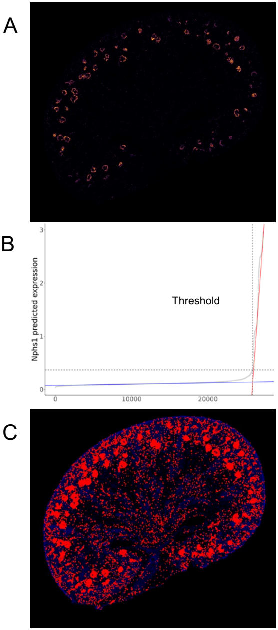
A
B
3
2
1
0
Nphs1 predicted expression
Threshold
0
10000
20000
C

Figure S11

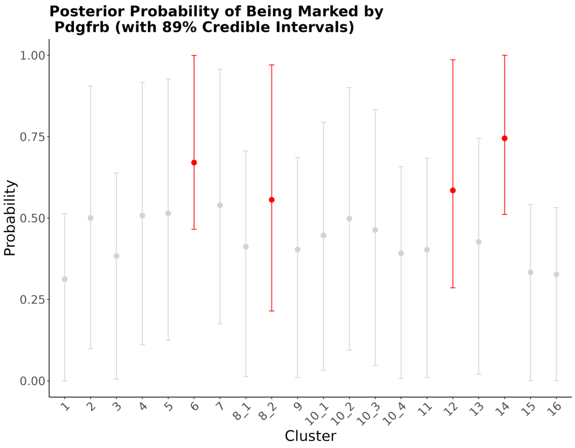
Posterior Probability of Being Marked by
Pdgfrb (with 89% Credible Intervals)
1.00
0.75
0.50
0.25
0.00
Probability
1
2
3
4
5
6
7
8_1
8_2
9
10_1
10_2
10_3
10_4
11
12
13
14
15
16
Cluster

Figure S1 Figure S1. Identification of Landmark Cell Types using Single-Cell Transcriptomics
(A) Successful integration of publicly available embryonic day 18.5 mouse kidney single-cell RNA-seq datasets (GEO accession numbers GSE108291 and GSE157594) is shown in the combined UMAP plot.
(B) Coarse annotation of the integrated dataset based on the expression of canonical marker genes, which are expected to be cell-type specific.

Figure S2. Landmarks measured by CARTANA.
Spatial localization of landmark genes measured by CARTANA (A, E15; B E18, C P3). The figure shows in-situ sequencing signal of all landmark genes, grouped according to their targeted cell type. Red markers indicate the spatial location of landmark genes on a DAPI-stained background image, using coordinates from CARTANA results.

Figure S3. Prediction of spatial expression at E15 and P3.
The predicted spatial distributions of Pou3f3, Ass1, Cldn11 and Apcdd1 (depicted at E18.5 in Figure 3) at the E15 (A-D) and P3 (E-H) stages indicating the extensibility of KSTAT.

Figure S4. Heterogeneous spatial expression of mural cell marker genes.
(A-D) EIGEN analysis predicted that the genes Mgp, Cspg4, Acta2 and Abcc9 would demonstrate relative specificity for the four clusters of putative mural cells identified in Figure 7. Here we present UMAP's of isolated interstitial cells colored by expression for these genes.
(E) The predicted spatial expression of these four genes is simultaneously projected onto the 2D E18.5 kidney section (Mgp, red; Cspg4, white; Acta2, orange; Abcc9, yellow).
(F-I) The individual predicted spatial expression for the four genes.
(J-K) The predicted spatial expression for the canonical mural cell markers Rgs5 and Pdgfrb are shown for comparison.
(L) A composite image of RNAscope measurement of mRNA expression for Mgp, Cspg4, Acta2 and Abcc9 in E18.5 kidney that is largely consistent with the predictions by KSTAT, but implies additional heterogeneity.
(M-P) Greyscale images of the individual RNAscope measurement channels for Mgp, Cspg4, Acta2 and Abcc9.
(Q) Greyscale image of the mRNA expression of endothelial marker Pecam1 measured with RNA scope to demonstrate the location of blood vessels in the E18.5 kidney.

Figure S5. Spatial heterogeneity of expression of soluble guanyl cyclases.
(A-E) Predicted spatial expression of the individual components of the heterodimeric soluble guanyl cyclases at E18.5 (A-C) and P3 (D-F). The coloring of the gene symbols corresponds to that of the points indicating predicted expression of the gene in Figure 5.

Figure S6. Relationship between proportion of cells expressing landmark and intercluster variability.
The four landmark genes with the highest total measured signal in the in situ sequencing experiment are characterized by a higher proportion of nuclei expressing the gene above a threshold relative to the intercluster variance of the gene's expression in the snRNA-seq dataset.

Figure S7. Transcriptomic dataset used for inference.
This single-nucleus RNA-seq dataset from e18.5 mouse kidney was used as input to KSTAT to infer gene expression at specific locations in the kidney. The UMAP plot shows the transcriptomic diversity of the dataset, with cells colored by annotated cell type to demonstrate that all expected kidney cell types are represented, including endothelium (End), interstitium (Int), leukocyte (Leu), nephron epithelium (NE), podocyte (Pod), and ureteric epithelium (UE)

Figure S8. Distributions of normalized and imputed expression values.

In panel A we show the distribution of normalized counts for the landmark Smoc2. Note that the y-axis is on a log-scale. This was necessary for visual clarity since the data is zero-inflated. In panel B we show distribution of expression values imputed with MAGIC. This data can be more reasonably modeled as a mixture of two Gaussian distributions.

Figure S9. Overview of the KSTAT algorithm.
The KSTAT algorithm integrates spatially resolved transcriptomics data with single-cell RNA sequencing data to infer gene expression at specific locations. The resolution of the reference space was decreased by using a bin size of 256 x 256 pixels for illustration purposes.
(A) In situ sequencing generates a binary map for each landmark gene, where pixels in the reference space are designated as "on" (black) if fluorescence is detected or "off" (white) otherwise. Concurrently, the measured expression of the landmark in the snRNA-seq dataset is modeled as a bimodal mixture of two normal distributions, representing cells that express the gene and those that do not.
(B) For each pixel, a specific combination of parameters is selected based on the fluorescence detection status for individual landmarks at that location. The components are combined to yield a multivariate normal distribution. By evaluating the probability density function at a cell's landmark expression profile, we obtain the likelihood that the cell is located at that bin. This can be efficiently represented as an nc x nb left-stochastic matrix, where nc is the number of cells and nb is the number of bins. The dot product of a gene's expression vector (row in the expression matrix) and a column from the probability matrix yields the expected value of expression for that gene at the corresponding bin.
(C) The calculated expected expression for a gene can be visualized as a three-dimensional surface. A plane parallel to the x-y plane, located at height t along the z-axis, separates regions with expected expression values of at least t, enabling the identification of specific locations with high expected expression.

Figure S10. Automatic Thresholding of Expected Gene Expression
(A) Expected expression of Nphs1 calculated using KSTAT and projected into the reference space.
(B) Sorted expected expression reveals an s-shaped curve, where the elbow point distinguishes bins with negligible expression from those containing significant signal. Two linear models are fit to the data, and their intersection point is used to determine the threshold value for filtering.
(C) The filtered image highlights the location of the most significant Nphs1 expression.

Figure S11. Probability of cluster being marked by Pdgfrb expression.
We computed the posterior probability that each cluster of interstitial cells would be marked by Pdgfrb expression. This plot depicts the 89% credible intervals for this probability for each cluster. The intervals for the four clusters with the highest posterior probability are colored red.

### Tables

#### Table S1. Reconstruction error for landmark genes.

| Landmark gene | Reconstruction error (full) | p-value | Reconstruction error (LOOCV) | p-value |
|---|---|---|---|---|
| Ace | 592.7803 | 0 | 794.9769 | 0 |
| Acta2 | 840.4386 | 0 | 1281.791 | 0 |
| Adamts2 | 1939.5471 | 0 | 4896.1777 | 0 |
| Adamts5 | 231.19473 | 0 | 315.5603 | 0 |
| Agtr1a | 419.86465 | 0 | 1015.5008 | 0 |
| Aldob | 596.4809 | 0 | 869.4812 | 0 |
| Apoe | 4.276144 | 0 | 334.21423 | 0 |
| Aqp1 | 2915.077 | 0 | 3034.2988 | 0 |
| Aqp3 | 29.063541 | 0 | 1629.1128 | 0 |
| Atp1b1 | 63.55609 | 0 | 245.17203 | 0 |
| Bmp3 | 65.98732 | 0 | 985.7909 | 0 |
| Bmper | 121.84595 | 0 | 2735.4553 | 0.0004 |
| C1qb | 1173.8735 | 0 | 1385.0999 | 0 |
| Calb1 | 89.899475 | 0 | 3965.7373 | 0 |
| Ccnd1 | 42.59207 | 0 | 282.76263 | 0 |
| Cdkn1c | 409.6261 | 0 | 4034.3135 | 0 |
| Cited1 | 1321.2715 | 0 | 1326.2123 | 0 |
| Clca3a1 | 267.19098 | 0 | 894.39014 | 0 |
| Cldn1 | 873.1045 | 0 | 574.28894 | 0 |
| Cldn19 | 424.38123 | 0 | 624.3763 | 0 |
| Col14a1 | 346.964 | 0 | 256.73468 | 0 |
| Col4a4 | 1389.5942 | 0 | 2569.4248 | 0 |
| Cpn1 | 1400.9858 | 0 | 1422.1536 | 0 |
| Crabp2 | 7615.647 | 0.9469 | 1902.7458 | 0 |
| Crlf1 | 3781.2483 | 0.0023 | 4123.222 | 0.0105 |
| Cyp2j5 | 115.4169 | 0 | 638.25903 | 0 |
| Dcn | 2053.769 | 0 | 4953.993 | 0 |
| Dkk1 | 487.45096 | 0 | 6825.018 | 0.0002 |
| Dkk2 | 7547.0713 | 1 | 2904.4326 | 0 |
| Edn1 | 2158.4153 | 0 | 847.7621 | 0 |
| Egfl7 | 4.412018 | 0 | 377.74078 | 0 |
| Egfl8 | 2017.7986 | 0 | 1977.4136 | 0 |
| Epcam | 285.5703 | 0 | 216.75189 | 0 |
| Fabp4 | 141.49905 | 0 | 143.84642 | 0 |
| Fibin | 2416.9724 | 0 | 4172.3022 | 0.0012 |
| Foxd1 | 224.92447 | 0 | 4997.4453 | 0.8156 |
| Fxyd2 | 130.90854 | 0 | 276.71716 | 0 |
| Fxyd3 | 363.2724 | 0 | 2576.2085 | 0 |
| Gata3 | 39.589386 | 0 | 582.03204 | 0 |
| Gm13889 | 896.47253 | 0 | 727.1924 | 0 |
| H19 | 66.72705 | 0 | 383.74414 | 0 |
| Irx1 | 73.47623 | 0 | 1409.5747 | 0 |
| Itga1 | 1617.7755 | 0 | 1346.5057 | 0 |
| Khdrbs3 | 523.4767 | 0 | 334.07495 | 0 |
| Krt18 | 121.275345 | 0 | 215.85541 | 0 |
| Krt19 | 512.81366 | 0 | 2845.2456 | 0 |
| Lcn2 | 2290.026 | 0 | 1460.5105 | 0 |
| Ldhb | 312.76855 | 0 | 751.4701 | 0 |
| Lox | 11.50905 | 0 | 119.16601 | 0 |
| Ly6a | 527.48303 | 0 | 671.5596 | 0 |
| Mafb | 802.4355 | 0 | 974.58575 | 0 |
| Mal | 857.7024 | 0 | 1184.7786 | 0 |
| Mgp | 913.16 | 0 | 889.43634 | 0 |
| Myh11 | 714.8053 | 0 | 15812.24 | 0 |
| Ndufa4l2 | 5871.611 | 0 | 3011.6877 | 0 |
| Nphs1 | 451.6538 | 0 | 771.3397 | 0 |
| Npnt | 550.5022 | 0 | 829.5585 | 0 |
| Nrp2 | 688.3566 | 0 | 729.10803 | 0 |
| Nts | 246.242 | 0 | 576.1212 | 0 |
| Pdzk1 | 606.09454 | 0 | 622.7868 | 0 |
| Penk | 39.309387 | 0 | 578.85425 | 0 |
| Podxl | 1.1750536 | 0 | 415.846 | 0 |
| Prrx1 | 2674.9304 | 0 | 1285.189 | 0 |
| Pttg1 | 3289.2952 | 0 | 2607.7266 | 0 |
| Rbp4 | 656.0679 | 0 | 756.3583 | 0 |
| Rgs5 | 1050.9075 | 0 | 2800.9683 | 0 |
| Rprm | 2956.7502 | 0.0005 | 10170.801 | 1 |
| S100a8 | 5917.1797 | 0.0008 | 7430.7344 | 0.0205 |
| S100g | 2439.9175 | 0 | 3053.4954 | 0 |
| Six2 | 412.72992 | 0 | 6960.844 | 0.1189 |
| Slc27a2 | 3.3423166 | 0 | 539.2528 | 0 |
| Slfn9 | 1587.6963 | 0 | 881.9777 | 0 |
| Smoc2 | 420.83698 | 0 | 1763.409 | 0 |
| Sox17 | 426.28735 | 0 | 761.8478 | 0 |
| Stmn2 | 318.62225 | 0 | 6015.0625 | 0.0436 |
| Sytl2 | 875.6852 | 0 | 765.98645 | 0 |
| Tagln | 308.05737 | 0 | 1722.0133 | 0 |
| Tfdp1 | 234.39389 | 0 | 283.59528 | 0 |
| Thbs2 | 63.78773 | 0 | 630.5244 | 0 |
| Timm8a1 | 2602.7458 | 0 | 992.02814 | 0 |
| Tm4sf1 | 774.6371 | 0 | 1640.4381 | 0 |
| Twist1 | 1269.4514 | 0 | 3564.3967 | 0.0002 |
| Tyms | 90.73059 | 0 | 195.61588 | 0 |
| Tyrobp | 8824.809 | 0.0067 | 20546.783 | 0.3577 |
| Wfdc2 | 1211.4172 | 0 | 1279.9431 | 0 |
| Wt1 | 7788.66 | 0.0002 | 7883.96 | 0.0007 |

#### Table S2. Ligand-receptor interacition scoring.

| CellPhoneDB | | | | | | KSTAT | | | | | |
|---|---|---|---|---|---|---|---|---|---|---|---|
| Interaction ID | Interactors | Rank | Mean | Score | p-value | Interaction ID | Interactors | Ligand mass (l) | Receptor mass (m) | Sinkhorn divergence (d) | Likelihood (L) | Percentile |
| CPI-SS0E41702C1 | IGF1-IGF1R | 0.5 | 1.141 | 100 | 0 | CPI-SC0456039E0 | RSPO1-KREMEN1+LRP6 | 126878.625 | 91675.54688 | 86069.96875 | 135142 | 100 |
| CPI-SS063D79C85 | DKK2-LRP6 | 0.5 | 0.695 | 100 | 0 | CPI-SC00DB2B867 | BMP4-ACVR2A+BMPR1B | 594907.125 | 97008.84375 | 653801.75 | 88270.25781 | 99.81343284 |
| CPI-SS0703C3CE4 | DKK1-LRP6 | 0.5 | 0.671 | 100 | 0 | CPI-SC052486D3E | BMP4-ACVR2A+BMPR1A | 594907.125 | 97008.83594 | 653801.875 | 88270.24219 | 99.62686567 |
| **CPI-SC0E149DD74** | **WNT5A-FZD4+LRP6** | **0.5** | **0.254** | **100** | **0** | CPI-SS0F81DBF22 | EFNA4-EPHA7 | 340176.75 | 114136.3906 | 727982.875 | 53334.42188 | 99.25373134 |
| CPI-SC04AF69F59 | WNT4-FZD4+LRP6 | 0.75 | 0.206 | 100 | 0 | CPI-CS09F74FD39 | GLS+SLC1A3-GRM7 | 267084.3438 | 251563.7656 | 1432009.5 | 46919.20313 | 99.06716418 |
| CPI-SC05E81C125 | WNT5A-FZD8+LRP6 | 0.5 | 0.173 | 100 | 0 | CPI-SS04A97448D | FGF1-FGFR2 | 107995.2266 | 373260.5625 | 883960.5 | 45601.98828 | 98.88059701 |
| CPI-SC01D1E05E8 | WNT5A-FZD6+LRP5 | 0.5 | 0.165 | 100 | 0 | CPI-SS0109FE6AE | HGF-MET | 262020.8906 | 94536.57031 | 660154.4375 | 37522.36328 | 98.69402985 |
| CPI-SC039EE3D74 | WNT5A-FZD6+LRP6 | 0.5 | 0.165 | 100 | 0 | CPI-SS00508D1C2 | VEGFB-NRP1 | 580193.375 | 241630.875 | 3962320 | 35381.44922 | 98.50746269 |
| CPI-SC027A59AFB | WNT5A-FZD5+LRP5 | 0.5 | 0.153 | 100 | 0 | CPI-SS09947DDD9 | EGF-EGFR | 326812.0625 | 34355.80469 | 322971.875 | 34764.30078 | 98.32089552 |
| CPI-SC0DD562B41 | WNT5A-FZD5+LRP6 | 0.5 | 0.153 | 100 | 0 | CPI-SS0659DBE72 | EFNA4-EPHA4 | 340176.75 | 149737.9688 | 1506252.75 | 33817.28516 | 98.13432836 |
| CPI-SC0B4243FE6 | WNT7B-FZD4+LRP6 | 0.5 | 0.146 | 100 | 0 | CPI-SC07CA7AF54 | DKK1-KREMEN1+LRP6 | 201819.7031 | 91675.54688 | 557737.75 | 33173.17188 | 97.94776119 |
| CPI-SC0B697A1AC | WNT9B-FZD4+LRP6 | 0.5 | 0.145 | 100 | 0 | CPI-SS021718DBF | TENM2-ADGRL2 | 279939.7188 | 307189.375 | 3043228.25 | 28257.6582 | 97.76119403 |
| CPI-SS093707FE0 | CLSTN2-NRXN2 | 0.75 | 0.124 | 100 | 0 | CPI-SS0BCB3D402 | TENM2-ADGRL3 | 279939.7188 | 289112.7188 | 2917999 | 27736.17578 | 97.57462687 |
| CPI-SC023CD2194 | WNT8B-FZD4+LRP6 | 0.5 | 0.122 | 100 | 0 | CPI-CS099F73A95 | SULT1A1-PPARA | 143263.5 | 50757.23438 | 285304.5625 | 25487.35547 | 97.3880597 |
| CPI-SS0BF9265F0 | CLSTN1-NRXN2 | 0.5 | 0.108 | 100 | 0 | CPI-SS06157E5AF | LRRTM4-NRXN2 | 128232.6641 | 42899.08594 | 223787.2344 | 24581.67188 | 97.20149254 |
| CPI-SS0934535FF | NLGN2-NRXN2 | 0.75 | 0.086 | 100 | 0 | CPI-CS0D89E7C19 | SULT1A1-PPARD | 143263.5 | 45404.95313 | 287083.5625 | 22658.46289 | 97.01492537 |
| CPI-SS06157E5AF | LRRTM4-NRXN2 | 0.25 | 0.084 | 100 | 0 | **CPI-SC00BDFD604** | **WNT9B-FZD2+LRP5** | **328724.7813** | **56320.08984** | **842544.5** | **21973.6875** | **96.45522388** |
| CPI-SS023A16921 | LRRTM2-NRXN2 | 0.5 | 0.069 | 100 | 0 | CPI-SC01C6EA11C | BMP4-ACVR2B+BMPR1B | 594907.125 | 60052.10547 | 1673202 | 21351.5332 | 96.08208955 |
| CPI-SS08D8F6957 | LRRTM3-NRXN2 | 0.5 | 0.067 | 100 | 0 | CPI-SC01D95B14C | BMP4-ACVR2B+BMPR1A | 594907.125 | 60052.10547 | 1673202 | 21351.5332 | 96.08208955 |
| CPI-CS0259A0EB4 | UBASH3B-PPARA | 0.5 | NA | 100 | 0 | CPI-SS0CFABB69A | FGF8-FGFR3 | 269302.5625 | 59363.04297 | 763375.75 | 20942.00586 | 95.52238806 |
